# Neuronal activity in the premotor cortex of monkeys reflects both cue salience and motivation for action generation and inhibition

**DOI:** 10.1101/796417

**Authors:** Margherita Giamundo, Franco Giarrocco, Emiliano Brunamonti, Francesco Fabbrini, Pierpaolo Pani, Stefano Ferraina

## Abstract

Animals adopt different strategies, promoting certain actions and withholding inconvenient ones, to achieve their goals. The motivation to obtain them is the main drive that determines the behavioural performance. While much work has focused on understanding how motor cortices control actions, their role on motivated behaviours remains unclear. We recorded from dorsal premotor cortex (PMd) of monkeys performing a modified version of the stop-signal task, in which the motivation to perform/withhold an action was manipulated by presenting cues that informed on the probability to obtain different amounts of reward in relation to the motor outcome. According to the motivational context, animals performance adapted to maximize reward. Neuronal activity displayed a cue salience related modulation at trial start and, while the behavioural response approached, reflected more the motivation to start/cancel the action. These findings reveal multiple representations of motivation-related signals in PMd, highlighting its involvement in the control of finalized actions.

**SIGNIFICATIVE STATEMENT:** The motivation to obtain rewards drives how animals act over their environment. To explore the involvement of motor cortices in motivated behaviours, we recorded high-resolution neuronal activity in the premotor cortex of monkeys performing a task that manipulated the motivation to generate/withhold a movement through different cued reward probabilities. Our results show the presence of neuronal signals dynamically reflecting a cue related activity, in the time immediately following its presentation, and a motivation related activity in performing (or cancelling) a motor program, while the behavioural response approached. The encoding of multiple reward-related signals in motor regions, leads to consider an important role of premotor areas in the reward circuitry.

## INTRODUCTION

To achieve the desired goals both humans and animals strategically adjust their behaviour. Optimality is often obtained by either acting on the environment or refraining programmed actions when no more convenient, accordingly to the specific context. In these situations, stimuli predicting reward become salient, independently of their physical properties, and motivate the subjects to prepare (or refrain) for action.

Electrophysiological, imaging and lesion studies succeeded in identifying motivation-related and reward-related signals in different brain areas^1–3^. While prefrontal and parietal cortex seem more involved in encoding the value and salience of the sensory signals used to inform about the incoming reward, the activity in motor cortical areas was proposed to be mostly related to the motivation to perform an action^4,5^ even if modulated by other factors like reward expectation^6,7^, reward feedback^8^, and anticipation of reward delivery^9^.

Being difficult to separate motivation from salience in tasks ending with a motor act and a linked reward, it is still unknown if the salience of the cue, used to inform on the reward context, it is encoded in motor areas. Moreover, it is still unknown if the motivation induced by the reward prospect could also influence the inhibitory control.

In this study we focused on the dorsal premotor cortex (PMd), a frontal motor area crucial for the formation of action plans^10,11^, for the executive control of motor acts, specifically motor inhibition^12–14^, and known to be modulated by the motivation to act^6^, through a multielectrode approach to study neuronal modulation in non-human primates.

To study both cue salience and motivation encoding in PMd, we combined a cueing paradigm with a well-established motor control task, the stop-signal task^15^, requiring often the completion (go trials) but sometime the cancellation (stop trials) of a movement when properly signalled. In the present version of the stop-signal task, a salient or a neutral cue was presented at the beginning of each trial, when the animals were still unaware of the type of response necessary to get the reward (moving or withholding). Two salient cues were designed to motivate one of the two possible motor outcomes and discourage the other one. In the Go+ condition the cue signalled that if the ongoing trial will end with a request of movement execution (go trials), the monkey would receive a higher amount of reward respect to the trials ending with a movement cancellation (stop trials). On the contrary, in the Stop+ condition the amount of reward between movement cancellation and execution was reversed. Lastly, a neutral (non-salient; control) cue condition was envisaged, in which the amount of reward was less than the highest possible and equally distributed either when the movement was successfully executed or cancelled.

Aim of this study was to disentangle the neuronal encoding of the cue salience, emerging at the time of cue presentation, from the encoding of motivation, emerging after the presentation of the go signal (or the the stop signal). By using this approach, we expected to observe: 1) a modulation of the performance aimed at maximizing the opportunity to obtain the reward depending on the cued context; 2) a different response of the neuronal activity for the salient cues respect to the neutral cue, in the time following their presentation; 3) a modulation of the neuronal activity reflecting the motivation to execute or cancel an action on the basis of the expected reward. According to the different experimental conditions, the encoding of reward expectation should entail the maximum degree of motivation in performing a movement in the Go+ condition, a medium degree in the neutral condition and the lower degree in the Stop+ condition. This level of motivation should be reflected in the pattern of movement related neuronal activity. On the contrary, the maximal motivation in halting a movement should be detected in the Stop+ condition, an intermediate degree in the neutral condition and a minimum level in the Go+ condition with the stopping related activity modulated accordingly.

In agreement with our hypothesis, we found that the animals adopted a strategy strongly influenced by the information provided by the sensory cue to maximize the chance of getting as more reward as possible. Furthermore, we observed that PMd neuronal activity encoded the cue (salience) in the time immediately following its presentation and that in PMd a motivation related activity in performing or cancelling a motor program emerges at the time around movement initiation or cancellation, respectively.

Taken together, our results suggest that PMd activity is involved in the control of motivated behaviours, necessary for the functional adaptation to the environment demands.

## RESULTS

### Monkeys strategically adjust their behaviour based on the motivational context

To investigate whether the manipulation of the motivational context, as triggered by reward prospect, affects the behavioural performance during both the execution and the inhibition of a movement, three monkeys were trained in a modified version of the stop-signal task (Fig. 1a, b).

**Fig. 1.**
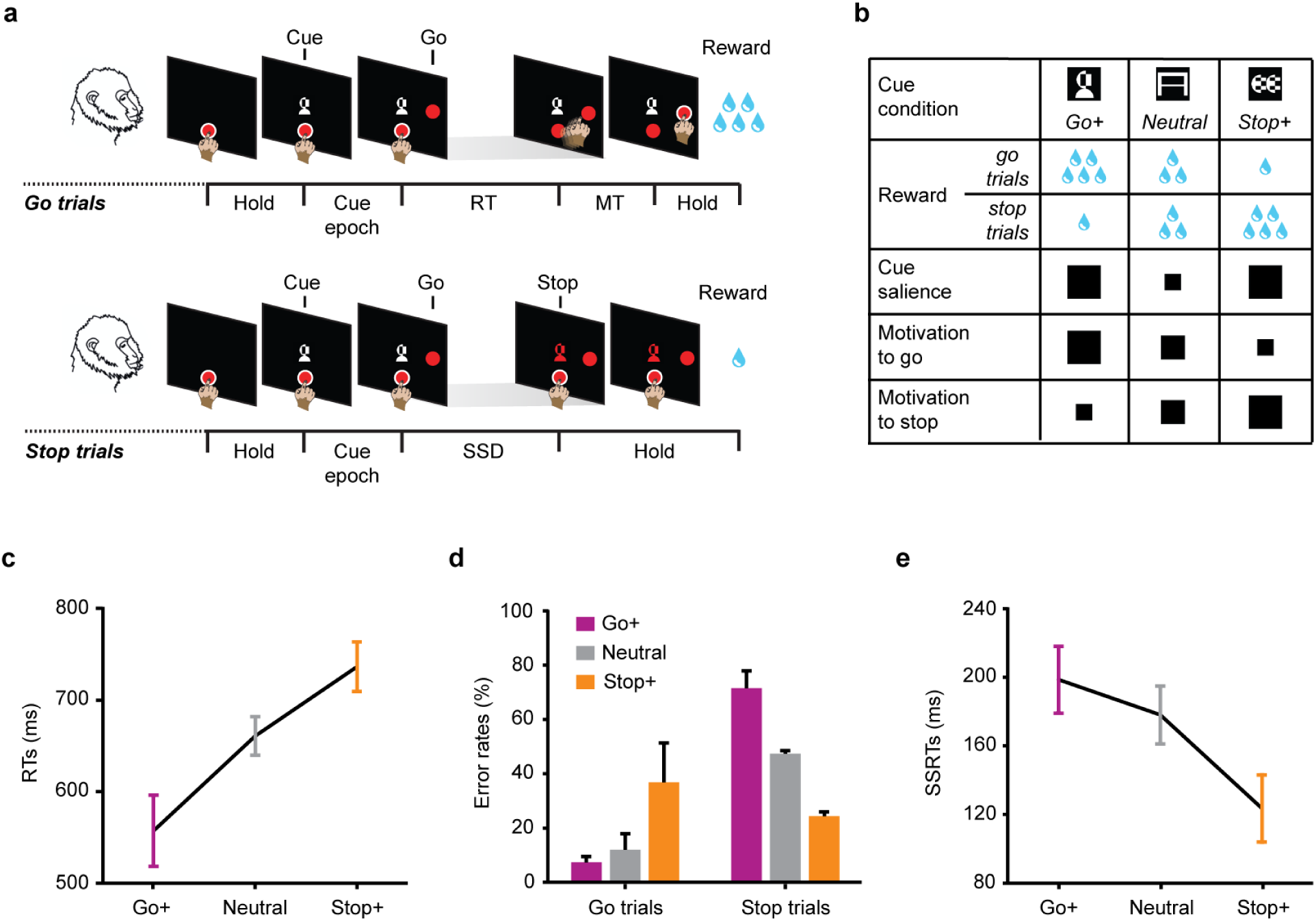
Motivational stop-signal task and behavioural performance. **a** Sequence of events for the Go+ condition. Each trial started when the monkeys touched a central target on the screen. After a variable delay, one of three possible cues (black and white abstract images; here the Go+ cue is indicated) appeared above the central target, informing about the amount of the expected reward. The monkeys were required to continue touching the central target (cue epoch). After 1000 ms, a peripheral target randomly appeared on the right or left of the screen. The peripheral target corresponded to a Go signal, instructing the monkeys to detach their hand from the central target (reaction time; RT) and to reach the peripheral target (movement time; MT) to receive the reward (go trials, 75%). Occasionally, at a variable delay from the Go signal (stop signal delay, SSD), the cue became red (Stop signal). In this instance, the monkeys had to hold the central target to earn the reward (stop trials, 25%). The white halo around either the central or the peripheral target was used as feedback of touch for the monkeys. Cue: cue presentation; Go: Go signal; Stop: Stop signal. **b** Schematic of the different levels of cue salience and motivation (either to go or to stop) for Go+, Neutral and Stop+ conditions. **c-e** Performance measures affected by motivational context. Reaction times (RTs) on correct go trials (mean ± SEM, n = 3 monkeys), percentage of errors (error rates) on go and stop trials (mean ± SEM, n = 3 monkeys) and estimated stop signal reaction times (SSRTs) (mean ± SEM, n = 12 sessions) were calculated for each cue condition.

Monkeys were instructed to reach a peripheral target (PT) after a Go signal (go trials), and to withhold their response (stop trials) when an unpredictable Stop signal followed after a variable delay (stop signal delay; SSD; see Methods for details). At the beginning of each trial, a visual cue appeared when the monkeys were still unaware of the type of trial they were performing, either go or stop. The cue informed about the amount of the expected reward in relation to the type of trial (go/stop). Three equiprobable and randomly intermingled cue conditions were envisaged: Go+ with larger reward for correct go trials compared to correct stop trials; Stop+ with smaller reward for correct go trials compared to correct stop trials; and Neutral control with equal amounts of reward for correct go and stop trials.

Manipulating the cue conditions in this way, we intended comparing two motivationally salient cues (Go+ and Stop+), after which we hypothesized a change in the behavioural strategy to maximize the probability to obtain the higher amount of reward, with a no salient Neutral cue condition, corresponding to the classical condition of stop-signal task, in which subjects simply have to perform correctly each of the trials, whether go or stop, to obtain the same amount of reward.

As expected, monkeys adopted different strategies based on the cue condition. We evaluated whether the amount of the expected reward influenced monkeys’ engagement in generating a movement, by analysing the duration of the reaction times (RT) and error rates (i.e. the probability of do not detach or to detach late) in go trials. We found that monkeys were faster in reacting to the Go signal and less prone to errors when a larger reward for go trials was at stake.

Overall, RTs were shorter for Go+ compared to Neutral and, in turn, Stop+ conditions (Fig. 1c; see also Supplementary Table 1 for further information from each monkey). This result achieved significance in all monkeys (mean ± SEM; monkey 1: Go+ 593 ± 4 ms; Neutral 675 ± 5 ms; Stop+ 786 ± 11 ms; monkey 2:

Go+ 480 ± 6 ms; Neutral 688 ± 7 ms; Stop+ 731 ± 9 ms; monkey 3: Go+ 599 ± 2 ms; Neutral 619 ± 3 ms; Stop+ 692 ± 4 ms; one-way ANOVA, all *p*’s < 0.0001). Furthermore, monkeys committed fewer errors in go trials on Go+ than on Neutral and, in turn, Stop+ condition (Fig. 1d). Examining each monkey separately (z score test), error rates were different in monkeys 2 and 3 across all possible comparisons (all *p*’s < 0.0001; monkey 2: Go+ 11%; Neutral 24%; Stop+ 41%; monkey 3: Go+ 3%; Neutral 6%; Stop+ 10%); whereas in monkey 1, error rates in Stop+ where higher compared to both Neutral and Go+ conditions (*p*’s < 0.0001), while no difference was found between these last two conditions (*p* = 0.503; Go+ 7%; Neutral 6%; Stop+ 59%).

We also evaluated whether the amount of the expected reward affected the inhibitory control. To this aim we compared the error rates in stop trials (i.e. probability of do not withhold the response), and the latency of the stop process (stop signal reaction time; SSRT; see Methods for details) across motivational contexts. We found that when the stopping was more rewarded the inhibitory control of the monkeys was more effective.

Particularly, monkeys committed about 50% of errors in the Neutral condition, as expected by the similarity here with the classical stop-signal paradigm (see Methods), whereas in the Stop+ and Go+ conditions respectively less and more errors were committed (Fig. 1d). The different inhibitory control for the three cue conditions was evident in each monkey across all possible comparisons (z score test, all *p*’s < 0.001; monkey 1: Go+ 69%; Neutral 50%; Stop+ 23%; monkey 2: Go+ 84%; Neutral 46%; Stop+ 22%; monkey 3: Go+ 62; Neutral 47%; Stop+ 28%). Lastly, the monkeys were faster in inhibiting their response when the stopping was more rewarded than the going; indeed, shorter SSRTs were observed in Stop+ condition than in Neutral and Go+ (Friedman test for repeated-measures, chi-square = 8.7917, *p* = 0.012; Fig. 1e).

Overall, these data show that all the monkeys adopted different behavioural strategies, producing shifts in the trade-off between responding and stopping based on motivational context (see also Supplementary Fig. 1).

### PMd neuronal activity signals both cue salience and motivation to move

We asked whether the observed behavioural effects could be tracked by the neuronal activity recorded from the dorsal premotor cortex (PMd; Supplementary Fig. 2). To this aim, we analysed the multiunit activity (MUA) extracted from 96-channel Utah arrays of one session for monkey 1, and two sessions for monkey 2, representing the best performance and adherence to the stop-signal paradigm (for details, see Table 1 and Methods). One-hundred-twenty-four recordings (80 from monkey 1 and 44 from monkey 2) were classified as task-related (see Methods) and used for the following analyses.

**Table 1.** Details of behavioural performance of sessions employed for the neuronal analyses.

| *Cue condition* |  | *Go+* | *Neutral* | *Stop+* |
| --- | --- | --- | --- | --- |
| ***Go trials*** |  |  |  |  |
| Error rate (%) | Monkey 1 | 7 | 4 | 43 |
|  | Monkey 2 | 9 | 21 | 35 |
| Mean RT ± SE (ms) | Monkey 1 | 569 ± 5 | 643 ± 7 | 792 ± 11 |
| | Monkey 2 | 433 ± 6 | 654 ± 8 | 717 ± 10 |
| ***Stop trials*** |  |  |  |  |
| Error rate (%) | Monkey 1 | 71 | 58 | 33 |
|  | Monkey 2 | 89 | 45 | 24 |
| Mean wrong-stop RT ± SE (ms) | Monkey 1 | 554 ± 10 | 607 ± 12 | 676 ± 25 |
| | Monkey 2 | 424 ± 11 | 569 ± 15 | 571 ± 27 |
| SSRT (ms) | Monkey 1 | 276 | 279 | 238 |
|  | Monkey 2 | 172 | 157 | 101 |

We first investigated whether PMd activity represents the degree of motivation to move, in direct association with behavioural performance, or other reward-related aspects. Indeed, from the cue presentation, which informs about the expected reward, the neuronal activity will possibly start to differentiate between cue conditions. We expected that if neuronal activity reflects the motivation to move, the recordings should follow the same pattern as the behavioural RTs: for example, in Go+ conditions, compared to the Neutral and Stop+, a higher level of activity should be observed, and this should be related to RT duration. Otherwise, neuronal activity could represent the cue salience: in this case, we expected higher neuronal activity slightly following motivationally salient cues – i.e. for Go+ and Stop+ - compared to Neutral condition.

To tackle these issues, we focused on the neuronal activity observed during go trials in the cue epoch of the task when no specific movement is needed to be yet prepared, and in the “Pre-MOV” epoch - i.e., a time after the Go signal where movement preparation should be on act (and where motor execution parameters like movement direction were observed; see Supplementary Fig. 3).

Fig. 2a shows two example recordings in which the neuronal modulation, related to the cue and to the movement preparation, changed over conditions and time. In some recordings (as for the example 1), the neuronal modulation started early after the cue presentation. Here the neuronal activity typically observed was higher for the Go+ and the Stop+ conditions than for the Neutral condition suggesting that what signalled was the salience of the cue. In the later epochs, the modulation reflected more the level of motivation to move, becoming higher for the Go+ than for Neutral and Stop+ conditions when approaching to the Pre-MOV epoch. In other recordings (as for the example 2), the difference in neuronal activity across conditions started during the cue epoch or during the Pre-MOV epoch, but immediately showed an increased modulation for the Go+ compared to Neutral and Stop+ conditions, suggesting more a code for motivation from start.

**Fig 2.**
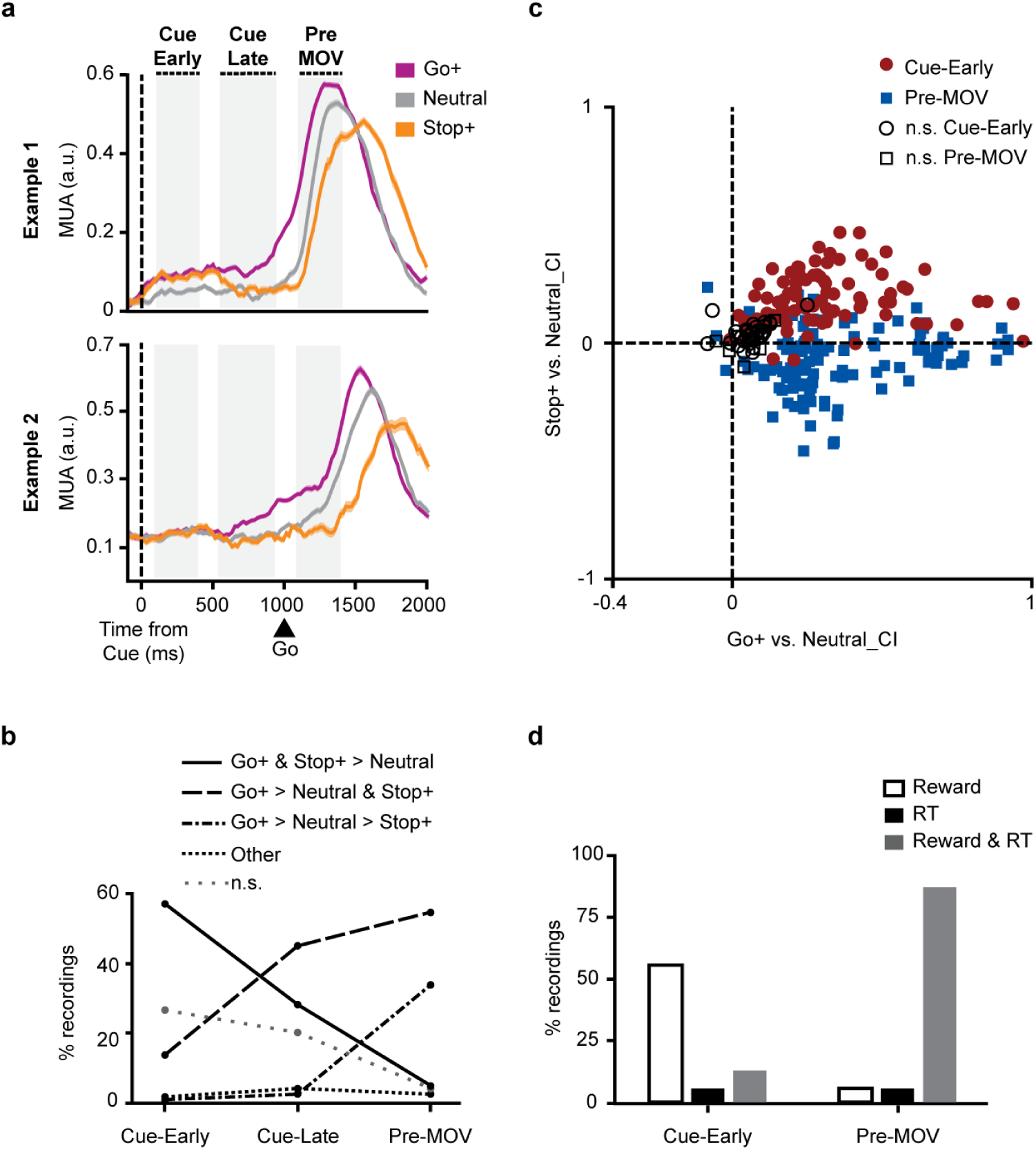
Effect of motivational context on the neuronal dynamics during go trials. **a** Average MUAs (± SE) from two example recordings are represented for each cue condition. Shaded grey areas depict the three trial epochs investigated in the following analyses: “Cue-Early” slightly following the cue presentation, “Cue-Late” just before the Go signal, and “Pre-MOV” following the Go signal. Go: Go signal. **b** Percentage of recordings belonging to each category based on their neuronal pattern across cue conditions (n = 124 recordings). Percentages were calculated separately for the three epochs. n.s.: not significant recordings. **c** Scatter plot of *Go+ vs. Neutral* and *Stop+ vs. Neutral* contrast indices (CI; n = 124 recordings). Contrast indices >0 indicate higher activity for salient cue conditions (Go+ or Stop+), and indices <0 indicate higher activity for the Neutral cue condition. Symbol color and style indicate the epoch in which indices were calculated. Black and white symbols represent those recordings with a not significant main effect of cue condition. **d** Percentage of recordings with a dependency of neuronal activity on reward-related factors (Reward), behavioural RTs (RT) or both (Reward & RT) (n = 124 recordings). Percentages were calculated separately for Cue-Early and Pre-MOV epochs.

To describe the occurrence of these patterns of modulation in our dataset, we classified recordings in 3 epochs of interest (Cue-Early, Cue-Late, Pre-MOV; see Methods) by a one-way ANOVA (with cue condition as factor, *p* < 0.05). Fig. 2b shows that-after the cue appearance - i.e. during the Cue-Early epoch - the activity in most task-related recordings was modulated more when salient cues were presented - i.e. for Go+ and Stop+ compared to Neutral condition (“Go+ & Stop+ > Neutral”: 57%); this pattern of activity became weakened during the Cue-Late epoch (28%), and consistently decreased during the Pre-MOV epoch (5%). Conversely, during Pre-MOV we observed higher levels of neuronal activity for Go+ than Neutral and Stop+ (“Go+ > Neutral & Stop+”: 55%, and “Go+ > Neutral > Stop+”: 34%). The different neuronal modulation observed from the Cue-Early epoch to the Pre-MOV (chi-square test: chi-square = 149.5309, *p* < 0.00001), suggests that PMd neuronal dynamics change across trial based on the motivational requests of the diverse task epochs.

To examine more in depth this topic, we calculated two contrast indices (CI) to emphasize the difference in activity between the Neutral and salient cue conditions (contrasting, Go+ versus Neutral, and Stop+ versus Neutral) during Cue-Early and Pre-MOV epochs. As illustrated in Fig. 2c, for most recordings, neuronal activity was higher for Go+ than Neutral in both epochs of the task, and this difference was stronger during Pre-MOV compared to Cue-Early epoch (Kruskal-Wallis test, chi-square = 15.09, *p* < 0.001); instead, the activity in Stop+ was higher than in the Neutral condition during Cue-Early epoch, and significantly inverted during Pre-MOV (Kruskal-Wallis test, chi-square = 122.39, *p* < 0.0001). Thus, it seems that, over the trial, neuronal activity in PMd first represents the salience of cues, then the motivation to perform the action when approaching to the initiation of the movement.

Finally, we investigated if the neuronal activity during the trial was better explained only by reward-related factors (salience during Cue-Early epoch and motivation to move during Pre-MOV), or it was better explained by movement preparation only, and so in association to behavioural RTs. To distinguish the motivation to move from movement preparation we considered the first one as a coarse measure (see Methods), defined by three degrees: high in Go+, medium in Neutral and low in Stop+. Otherwise, we hypothesized a finer relationship between movement preparation and RTs: the higher the movement preparation, the lower the RT.

Following the analysis described by Roesch and Olson^16^, we performed, for each recording, a multiple least-squares regression approach to calculate the parameters of 3 models representing the neuronal activity as a linear function of RTs and reward-related factors: 1) a reduced model incorporating RT only, 2) a reduced model incorporating reward-related factors only, and 3) a full model incorporating both factors. For each recording, we compared each of the reduced models to the full model using a nested F-test (see Methods). Fig. 2d shows the percentage of recordings with a significant improvement of fit only when the reward-related variable was considered in the model (white bar), only when RT variable was considered (black bar), or when they were both considered (grey bar). The results are in line with the data presented above, confirming that during Cue-Early epoch the neuronal activity of most recordings was mainly explained by reward-related factors, specifically salience (56%, 69 of 124); however, during the Pre-MOV epoch, the reward-related and RT-related effects were strongly yoked (87%, 108 of 124), suggesting that the neuronal activity explained by motivation is strictly related to motivation (behavioural RTs).

### In PMd, neuronal modulation during movement cancellation is influenced by motivation

We wanted also to investigate how neuronal activity in PMd is affected by the context change, as defined by the cue, during movement suppression. To this aim, we selected recordings showing a role in the inhibition process - i.e. their neuronal activity changed early enough in successful stop trials (after the Stop signal presentation but before the end of SSRT) to be able to influence the cancellation of the movement (see Methods for more details). We found that 85% (106 of 124, of which 75 from monkey 1 and 31 from monkey 2) of task-related recordings were differently modulated when the movement was inhibited compared to when it was performed. We classified them as countermanding (or CMT) recordings (see Supplementary Fig. 4).

Fig. 3a shows the neuronal modulation from three representative CMT recordings differently modulated by the motivational context: for all the examples, the left, central and right plots show the neuronal activity of correct stop trials (colour lines) and latency matched go trials (black lines), i.e., trials that have a similar level of movement preparation of the corresponding correct stop trials (see Methods) for Go+, Neutral and Stop+ conditions, respectively. Different patterns of modulation were observed. Indeed, we found that for most of recordings the activity decreased after the Stop signal presentation, and before the SSRT, in each cue condition, employing shorter times to suppress the neuronal activity when a large reward for stop trials was at stake (Stop+ condition) than under the Neutral condition and in turn the Go+ condition (example 1).

**Fig. 3.**
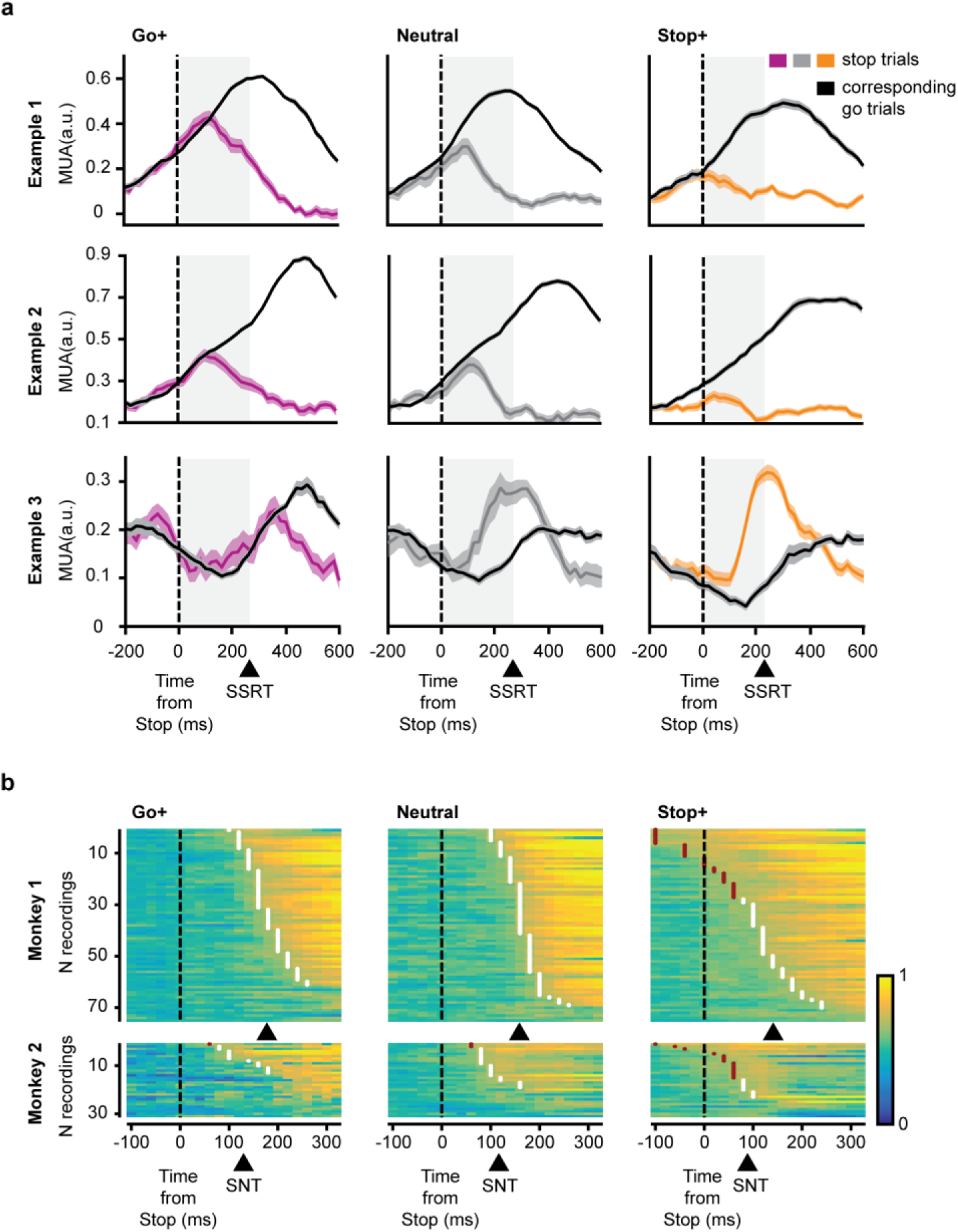
Different neuronal patterns for go and stop trials based on motivational context. **a** Three example recordings differently modulated by the motivational context. Left, central and right panels represent the neuronal activity for Go+, Neutral and Stop+ conditions, respectively. Each panel shows the average MUA (± SE) of correct stop trials and latency matched go trials. Shaded grey areas depict the duration of the stop signal reaction time (SSRT). **b** Time evolution of accuracy (auROC) in discriminating between correct stop trials and latency matched go trials (color plots). For each recording, the stop neuronal time (SNT) - i.e. the time at which the accuracy value reaches 0.65 – is depicted by white dots. Red dots represent unconsidered SNTs (either for the too short latency, <70 ms, or because preceding the Stop signal. The mean SNT for each monkey and condition is indicated by the black arrow below each panel.

However, for some recordings, the neuronal activity for the Stop+ condition was maintained low until the Stop signal presentation (example 2); in fact, in this instance, monkeys were more motivated to delay or not generate a movement to obtain the maximum reward amount. This aspect further suggests that the motivational context indicated by the cue was able to influence (proactively) the control of movement generation.

Lastly, we observed neuronal modulation examples where activity for the Go+ condition was not modulated or was modulated after the SSRT (example 3), compared to the other conditions. Together, these examples suggest that the motivation to inhibit a movement affects the time necessary to suppress the neuronal activity (stop neuronal time – SNT – i.e. the time of divergence between latency matched go trials and correct stop trials; see Methods), confirming what has been observed at the level of behavioural performance for SSRTs.

To find quantitative support to these suggestions, we performed a ROC analysis to identify, for each cue condition, the recordings involved in movement cancellation and to measure the SNT at the population level. Each plot in Fig. 3b, shows a measure of accuracy (ROC values) of the divergence of the neuronal activity obtained between latency matched go and correct stop trials, for all CMT recordings and cue conditions (Go+, Neutral, Stop+, respectively). Data are sorted by time at which the accuracy value reaches the threshold (ROC = 0.65) from the presentation of Stop signal.

We found that the SNT and the number of CMT-related recordings varied based on the motivational context. Particularly, 70% (74 of 106) of recordings revealed a divergent activity for Go+ condition; this population of recordings significantly increased to 83% (88 of 106) for Neutral (z score test; Go+ vs.

Neutral: z = −2.2649, *p* = 0.02382), and achieved the 88% (93 of 106) for Stop+ (Go+ vs. Stop+: z = - 3.1912, *p* = 0.00142); no difference was found between Neutral and Stop+ (Neutral vs. Stop+: z = - 0.9719, *p* = 0.33204). In Stop+ condition, the divergence between latency matched go and correct stop trials was evident even before or slightly after the Stop signal presentation (as observed in the example 2 of Fig. 3a): 40% (42 of 106) of the recordings distinguished between go and stop trials before or within 70 ms following the Stop signal (taking into account the expected latency for the information of the Stop signal to reach frontal lobe); whereas, in Neutral and Go+ conditions only 2 and 1 recordings were modulated at least at 60 ms after the Stop signal. Lastly, we found that the SNT was longer in Go+ (mean ± SE: 169 ± 5 ms) than in Neutral (152 ± 4 ms) and, in turn, Stop+ (135 ± 6 ms) conditions (one-way ANOVA; F(2,207) = 9.6429, *p* = 0.0001) (see Fig. 3b for SNTs calculated from each monkey).

To investigate more in depth how the motivational context affects the neuronal dynamics during the reactive inhibition process, we selected, for each cue condition, only recordings showing a reactive neuronal suppression pattern (at least 70 ms after the Stop signal presentation). For each recording, we calculated the time at which the neuronal activity in correct stop trials began to show a decreasing trend (decrease onset time), and the corresponding level of neuronal activity. Starting from this time we also computed the slope that characterizes the neuronal suppression pattern. We found clear differences in the neuronal dynamics based on the motivational context (see Fig. 4a for the modulation in one example recording). When the monkeys were more motivated to withhold a movement (Stop+ condition), the suppression of neuronal activity started slightly after the Stop signal presentation; differently, it was delayed for the Neutral and the Go+ conditions. Furthermore, in Stop+ condition, the mean neuronal activity was just slightly increased at the decrease onset time, and the following slope was smaller (Fig. 4b, c). When the preparation of the movement was more advanced - i.e. higher levels of neuronal activity at the decrease onset time - as in Neutral and Go+ conditions, a steeper suppression occurred (i.e. larger slopes) when the monkeys were more motivated to inhibit - i.e. in the Neutral compared to the Go+ condition.

**Fig. 4.**
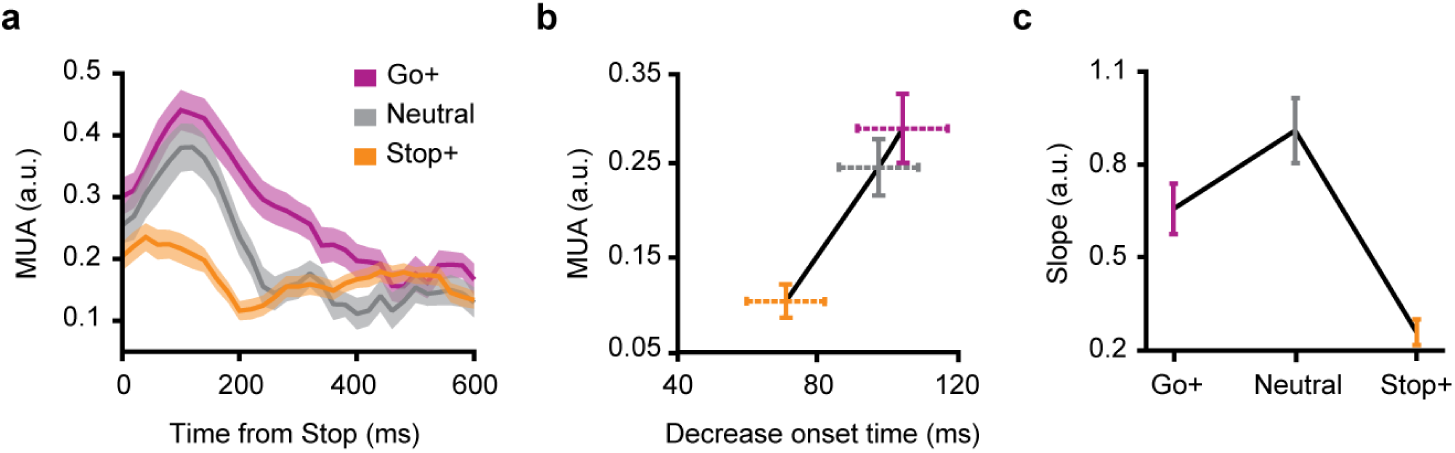
Neuronal modulation in stop trials based on the motivational context. **a** Example recording representing the average MUAs (± SE) in correct stop trials, for each cue condition. **b-c** Average (± SE) decrease onset time, the corresponding average MUA (± SE) and slope (± SE) of the neuronal modulations, calculated for each cue condition (n = 66 recordings for Go+, 75 for Neutral and 41 for Stop+).

Statistical testing supported the phenomenological pattern of neuronal activities: three separate one-way ANOVAs - one for the decrease onset times, the other for the corresponding neuronal activity, and the last one for the slopes, with factor cue condition - were done. The analysis of the decrease onset times showed that the neuronal activity was suppressed earlier in Stop+ condition, than in Neutral and Go+ conditions (F(2,179) = 4.5806, *p* = 0.01148); however, no significant difference between Neutral and Go+ conditions was found (Newman-Keuls post-hoc test: *p* = 0.518). Accordingly, the neuronal activity calculated at the decrease onset time was lower for Stop+, compared to Neutral and Go+ (F(2,179) = 14.064, *p* = 0.00000), but no difference was found between these last two conditions (Newman-Keuls post-hoc test: *p* = 0.217). The slopes too changed depending on the cue condition (F(2,179) = 14.863, *p* = 0.00000), but in a different manner: the Neutral condition showed a steeper slope compared to Go+ and, in turn, Stop+ (Newman-Keuls post-hoc test: all *p*’s < 0.05).

Thus, in general, we found that the higher the motivation to suppress a movement, the earlier the modulation of neuronal activity started and the lower the growth of activity was. Interestingly, for similar levels of neuronal activity at the decrease onset time, the higher motivation to inhibit a movement is translated in a stronger modulation (i.e. steeper slopes) after the Stop signal presentation.

### Population coding of salience and motivation in PMd

Once we found that PMd neuronal activity reflects both cue salience and motivation to act or to inhibit, we asked whether and how the information about the salience and the motivation is coded at the population level. To this aim we performed a neural decoding analysis by training a classifier to discriminate between the different conditions of the task using the whole population of task related recordings. We considered the salient cue conditions (Go+ and Stop+) with the Neutral one for “cue salience” and the three different cue conditions of the task for “motivation” (Go+ vs. Neutral vs. Stop+), including both go and correct-stop trials. Indeed, in both trial types information about salience and motivation must be represented in a similar way until the presentation of the Stop signal. A support to this logic of trials selection comes from performing a decoding analysis based on the trial type (go vs correct-stop) that shows the inability of the analysis to distinguish between these trial types until 200ms after the go signal (supplementary Fig. 5).

Figure 5 (white line in a and b) shows that PMd population activity encodes (classification accuracy; scale in the right side of a and b) the information about the “cue salience” just after the cue appearance (from 80 ms after the cue), and that this coding is kept almost constant above chance level (50%) over time. The encoding of the motivation (threshold 33%) is delayed (from 220 ms after the cue) compared to the encoding of the salience. However the strength of this last representation increases over time. These data suggest that at the population level neurons code for both salience and motivational signals but with different dynamics.

**Figure 5.**
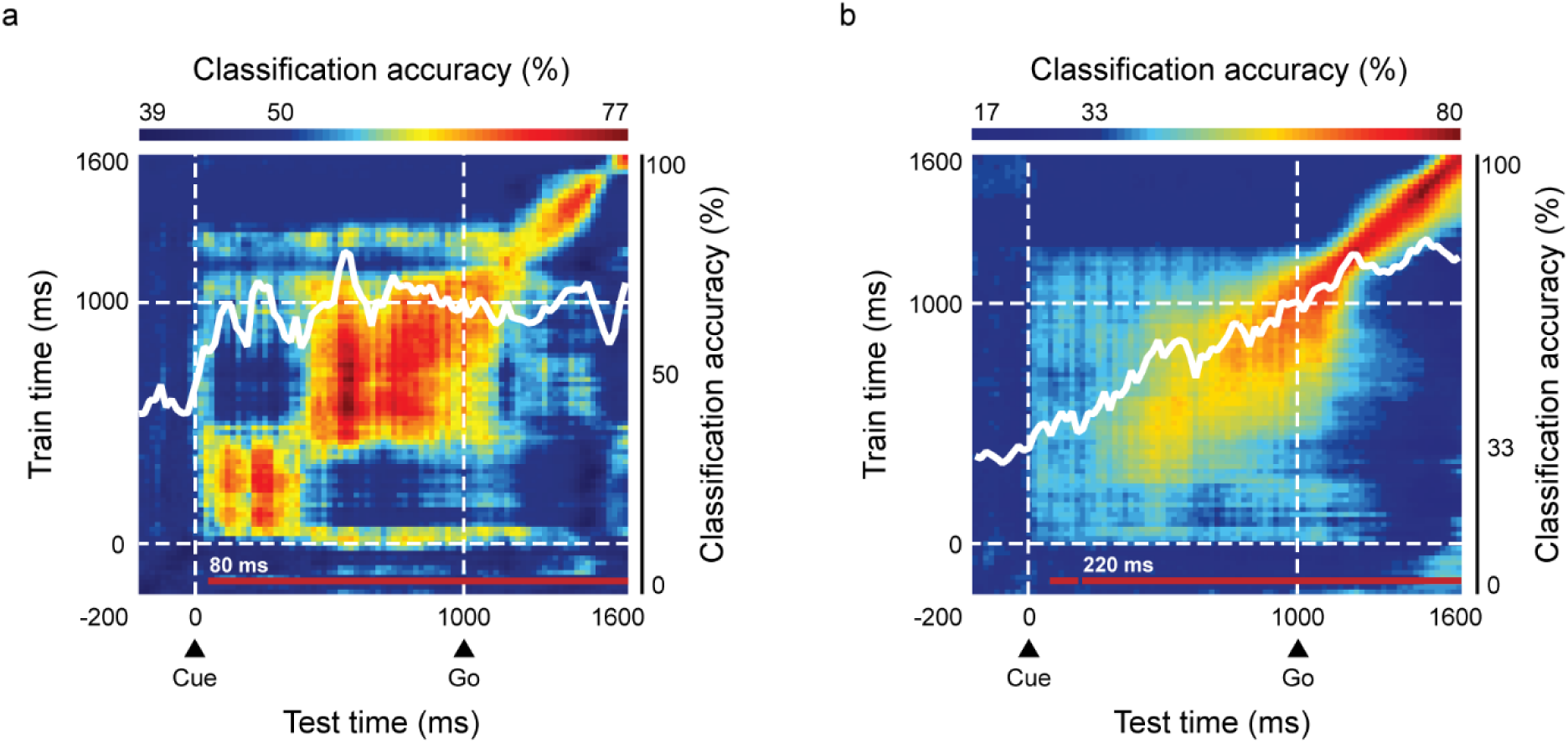
Classification accuracy over time and cross-temporal decoding plot for “cue salience” and “motivation”. The decoding analysis has been conducted to test separately the coding either of the cue salience (a) or the motivation (b) at the population level and to investigate the static/dynamic nature of the population coding. White lines represent the classification accuracy over time relative to the cue onset and to the go signal. Red lines in the lower part denote the times at which classification accuracy is above chance level (50% for salience and 33% for motivation, permutation test P < 0.05). Cross-temporal decoding plot: training and testing the classifier at different time points have tested accuracy of the classifier. A population coding it is considered to be dynamic if the accuracy decays rapidly when testing the decoder at different time points of those employed in the training phase (presence of a diagonal band in the plot); conversely it is considered to be static if the accuracy is preserved across different time points (presence of a square region in the plot).

To gain more insights on the nature of the representation of information for the different comparisons we performed a cross-temporal decoding analysis. Figure 5a shows that the static representation of the salience has two different phases (squares in the colour plot) in the early and late part of the cue epoch, respectively. Following the go signal the diagonal band is indicative of a more dynamic code, purportedly related to the bell shaped patterns of MUA before movement generation and the difference of RTs between conditions. When looking at the motivation signal we found a different pattern: a static code is less apparent in the cue epoch while a dynamic code emerges in the late part of the cue epoch (after about 300 ms from the cue onset) becoming more evident after the go-signal. Indeed around this point the motivation is translated in the proper level of movement preparation as driven by the cue.

Thus PMd neuronal activity is characterized by the representation of a salience signal after the cue and throughout the trial. After about 200 ms also the motivation is represented but as a growing signal that is translated in the proper level of movement preparation before the go signal, as observed also at the single recording level. Importantly for the second half of the cue-period these two signal coexists. This coexistence is further confirmed by the decoding analysis performed on correct-stop and go trials, where no difference between the two trial types can be detected until the go signal (supplementary Fig. 5).

## DISCUSSION

In this study, we intended to investigate which reward-related signals PMd encoded during both movement generation and inhibition. With this purpose, we presented a variant of the stop-signal task manipulating reward amounts in a novel way, to vary the motivational context. Our results suggest that PMd neuronal signals, representing the salience of cues, coexist with signals reflecting the motivation to move and to inhibit.

At the behavioural level, we observed that monkeys chose between different behavioural strategies to maximize the probability to obtain the higher reward amount. Particularly, when they were more motivated to move because of the higher reward for go trials, they committed more errors in stop trials, and responded more rapidly in go trials. Conversely, when stopping was prioritized, they were more accurate on stop trials and responded more slowly in go trials. This is in line with preceding findings that reveal the effects of motivational bias on stop-signal task performance^17,18^. Indeed, motivational bias can affect the decisional process about whether and when to initiate a response: the probability of responding and withholding will depend on the relative importance of the two goals. Confirmations come from human works on movement planning under risk, in which subjects are generally found to be very good at choosing motor strategies maximizing expected gain, when full information about the stimulus configuration and the assigned rewards is provided prior to movement onset^19,20^.

Neuronal correlates of behavioural performance were investigated during the period from the cue presentation to movement planning. We found that PMd neuronal activity after the cue presentation was differently modulated for Go+ and Stop+ (i.e., when the cue was informative of a possible high amount of reward associate to go or stop trials) respect to Neutral condition. We hypothesize that the two cues associated to higher reward amounts became salient cues for the monkeys. Indeed, these cues lead the monkeys to change their behavioural strategy to maximize the probability to obtain the higher reward. We interpreted this activity as encoding the salience of cues compared to other interrelated aspects of reward processing, such as the value of expected rewards or motivation.

The value, intended as the relative anticipated worth that some cue predicts^5^, is determined by the kind of reward (or penalty), its magnitude and probability^21^. Based on this definition, in our experimental paradigm, the value of the expected reward should be greater when monkeys expect higher reward for go trials (go trials representing the majority of trials presented to monkeys (75%) compared to stop trials (25%)), decreasing in the Neutral condition and, in turn, the condition associated to higher reward for stop trials. This is not consistent with our neuronal results observed after cue presentation. Indeed, the modulation was very high, not only for Go+ condition, but also for Stop+ despite the low occurrence of stop trials, and so the lower probability to obtain the higher reward in this condition. Accordingly, previous studies have shown that value of the expected reward is not represented by PMd neurons but is better explained by activity in prefrontal areas, such as OFC^6,16^.

Much more difficult is to distinguish between salience and motivation at the neural level. Although the two are intertwined, a stimulus is considered as “salient” if it leads to a general increase in arousal or is attention grabbing, whereas something is “motivational” if it enhances motor behaviours^5^. Furthermore, it has been suggested that signals related to motivation should be observed in the period leading up the behavioural response, whereas signals related to salience might solely be during the presentation of cues^5^. A previous finding supposes that PMd represents the motivation to act^6^. However, the experimental designs of this^6^ and following studies (for example ^22–24^) manipulate appetitive and aversive stimuli to dissociate value signals from signals related to motivation and salience; whereas they can’t demonstrate a distinction between these last two factors, since they covary in these tasks.

The task we designed may represent a novel experimental paradigm to better investigate this distinction and the brain areas involved. In fact, in this task the neuronal activity explained by salient cues doesn’t covary with the activity explained by the motivation to perform/suppress an action (for a similar approach, see Lin and Nicolelis^23^), allowing us to conclude that the early neuronal pattern of modulation, observed after the cue, encodes the salience. This pattern of activity changed into the time, becoming sensible to the degree of motivation when monkeys were preparing their behavioural response: the neuronal activity increased more when a large reward was expected for go trials, compared to the neutral condition and the condition associated to higher reward for stop trials, thus paralleling the RTs.

The dynamics is confirmed at the population level by means of the neuronal decoding analysis. Following the cue the population activity encodes the salience of the signal in the early period. After that the encoding of the salience continues in the late period when, at the same time, also the motivation starts to be encoded. In the late period of the cue epoch and towards the movement generation the neuronal activity codifies a specific level of movement preparation reflecting the motivation to move/refrain.

Accordingly, PMd represents not only the motivation to move but also the motivation to suppress actions, producing specific patterns of motor inhibition. After the presentation of a Stop signal, the neuronal activity decreased early when higher reward was at stake for stop trials, compared to the Neutral condition and, in turn, the condition associated to higher reward for go trials. These data suggest a “faster” stop-related decision process when monkeys are more motivated to suppress actions, as also confirmed by behaviour (SSRTs duration).

It could be assumed that the difference in inhibition patterns due to the different reward prospects is exclusively related to preparatory proactive processes allowing for differential preparation based on motivational cues^17,25,26^. However, we also suppose that the higher motivation to inhibit elicits, after the Stop signal presentation, enhanced reactive control mechanisms. In fact, for similar levels of movement preparation, the higher motivation to inhibit a movement was translated into steeper slopes after the Stop signal presentation.

Support to our observation comes from electroencephalography (EEG) and imaging studies. These studies designed a stop-signal task in which the expected reward was indicated by the color of the Stop signal itself, confirming that different reward prospects can influence reactive response inhibition without the involvement of global preparatory functions^27,28^.

All together, our neuronal results may be interpreted in the framework of decision-theoretic models (for review, see ^29–31^). These models suggest that when animals can choose between different potential actions, the variation in the expected reward exerts a correlated influence on both the choice behaviour of the animal and the neuronal activation in some motor-related areas^32–34^. The activity in these regions reflects both the action selection and movement preparation^35–37^.

In this framework, Pastor-Bernier and Cisek^32^ have conducted a study in which a monkey was trained to choose between two reach targets differently associated to reward values. The results showed that PMd activity was modulated by the relative value of potential reach targets, thus suggesting that decisions between actions are determined by a competition between action representations which takes place within sensorimotor circuits^32^.

Nevertheless, this idea has mainly been tested for spatial target selection. Our experimental design doesn’t allow to explicitly discriminate between activity patterns associated to spatial coordinates that correlate with different potential actions. Although we cannot discriminate between activity patterns associated to different potential actions, we can suppose that PMd may first represent decision-related variables such as the level of evidence in favour of a given behavioural choice: after the cue, PMd was more activated when the monkeys preferred a behavioural strategy (if move or withhold a movement in the current trial), depending on the expected reward amount; whereas, later in the trial, the activity reliably represented the monkey’s response choice.

## METHODS

### Animals

Three adult male rhesus monkeys (*Macaca mulatta*), 10-13 kg in weight, participated in this study. Monkeys had free access to food and controlled access to water during the experiments. They received fruit juice as a reward for performing the task. Animal care, housing and experimental procedures conformed to the European (Directive 210/63/EU) and Italian (DD.LL. 116/92 and 26/14) laws on the use of non-human primates in scientific research.

### Surgery

A single 96-channel Utah array (BlackRock Microsystem, USA) was implanted over the left PMd of monkey 1 and the right PMd of monkey 2 (using as anatomical landmarks after dura opening the arcuate sulcus and the pre-central dimple). The site of the implant was contralateral to the arm used during the experiment. The position of the chronic arrays of both monkeys is shown in an illustrative brain figure (Supplementary Fig. 2).

All the surgeries were performed under sterile conditions and veterinary supervision. Antibiotics and analgesics were administered postoperatively. Anesthesia was induced with ketamine (Imalgene, 10 mg kg−1 i.m.) and medetomidine hydrochloride (Domitor, 0.04 mg kg−1 i.m. or s.c.) and maintained by inhalate isoflurane (0.5–4%) in oxygen. Antibiotics were administered prophylactically during surgery and postoperatively for at least 1 week. Postoperative analgesics were given at least twice per day. Recordings started well after recovery from surgery (after a minimum of 10 weeks). A head-holding device was implanted in monkeys 2 and 3 before training started, while in monkey 1 the head-holder was implanted simultaneously with the array.

### Behavioural task

All the monkeys were trained to perform a motivational version of the stop-signal task intended to modulate the motivation to stop and to move by conditionally manipulating the reward amount. The monkeys were placed in a darkened, sound-attenuated chamber and seated in a primate chair with the head restrained facing a touch screen monitor (MicroTouch, 19 inches, 800 X 600 pixels resolution) 20 cm away. Liquid reward was delivered from a tube positioned between the monkeys’ lips, and eye movements were monitored by using a non-invasive Eye-tracker (Arrington Research Inc, AZ). The behavioural task was implemented using the software package Cortex (nimh.nih.gov) to control visual stimuli presentation and reward delivery, and to detect touches on the screen.

The task (Fig. 1a, b) consisted on go (75%) and stop trials (25%) randomly alternated. Each trial started when the monkeys touched a red central target (CT) on the screen. After a variable waiting time (400 ± 20 ms), a visual white cue appeared just slightly above the CT, informing about the amount of the expected reward. The monkeys were required to continue touching the CT for 1000 ms (cue epoch), after which a peripheral target (red circle; PT) randomly appeared in one out of two possible locations (i.e., at the right or left of the screen vertical midline). The PT presentation corresponded to a Go signal, instructing the monkeys to detach their hand from the CT (maximum time: 1400 ms for monkey 1, and 1200 ms for monkeys 2 and 3; reaction time, RT), and to reach the PT (movement time, MT). The monkeys had to touch it for a random time (900-1200 ms), until the reward was given (correct go trials). Occasionally, at a variable delay from the Go signal (stop signal delay, SSD), the cue became red (Stop signal) and, in this instance, the monkeys had to hold the central position for an additional interval (900-1000 ms interval) to earn the juice (correct stop trials). If the monkeys moved the hand from the CT, the trial was considered a wrong stop trial and no reward was given. Both correct and incorrect trials were followed by a 2000 ms inter-trial interval, during which the screen became black.

Within a session, three different cue conditions were selected with equal probability and intermixed in a random trial-by-trial fashion. The visual cue presented at the beginning of each trial indicated which of the three conditions was selected. Cues consisted in black and white abstract images (bitmaps, 16° x 16° of visual angle). The amount of reward for every cue condition was differently associated to go and stop trials as follows: Go+ condition was associated to a higher reward for successful go trials compared to successful stop trials (e.g. 5 vs. 1 juice drops); Stop+ condition was associated to reversed reward amounts (e.g. 1 vs. 5 juice drops); lastly, a Neutral control condition was associated with equal amounts of reward delivered for successful go and stop trials (3 vs. 3 juice drops).

In stop trials, SSDs were presented according to a fixed SSD procedure. Four progressively longer SSDs were computed so that, in the Neutral condition, the monkeys were able to successfully inhibit a movement in about 85%, 65%, 35%, and 15% (and, overall, in about 50%) of the stop trials (see ^12^ for a similar procedure). The same SSDs were employed for each of the cue conditions to compare stop performance and measures of inhibitory control based on shifts in motivational context.

### Behavioural analysis

To investigate the influence of motivational context on stop-signal task, we employed behavioural data from different sessions for three monkeys (monkey 1: 2 sessions; monkey 2: 3 sessions; monkey 3: 9 sessions). Data were processed using Matlab software (The MathWorks Inc., Natic, MA, USA) and statistical comparisons were performed by mean of the software Statistica (StatSoft, Inc.).

We tested error rates in go and stop trials (i.e. the probability of do not detach or to detach late in go trials, and the probability to detach in stop trials), merging data from different recording sessions, and calculating z-scores test (*p* < 0.05) from each pair of cue conditions, separately for each monkey.

To compare reaction times (RTs) among the three cue conditions, data collected in the different sessions were merged; we then performed a one-way ANOVA (*p* < 0.05) separately for each monkey.

Performance in stop trials was evaluated within the framework of the race model^15^, by calculating the latency of the stop process - i.e. the stop signal reaction time (SSRT).

The race model assumes that during stop trials two stochastic processes race toward a threshold: the go and the stop processes, triggered by the appearance of the Go signal and the Stop signal, respectively. The result of this race, either movement generation in wrong stop trials or movement inhibition in correct stop trials, will depend on which of these processes will reach its own threshold first. In correct stop trials, the stop process wins over the go process, and vice versa in wrong stop trials. The main assumption underlying the race model is that the go and stop processes are independent to each other (independence assumption). In particular, the model assumes two types of independence: first, that on a given trial, the latency of the go process does not depend on the latency of the stop process (stochastic independence); second, that the go process in stop trials must be the same as in go trials since the go process must be unaffected by the presence of the Stop signal (context independence).

Only if the independence assumption is validated, the race model could be used for the estimation of the SSRT. To validate the independence assumption, wrong stop trials RTs must be shorter than the correct go trials RTs^38^. We decided to calculate the SSRT only if the independence assumption was validated at least in the Neutral condition, i.e. when the amount of the expected reward was the same for correct stop and correct go trials (monkey 1: 2 sessions; monkey 2: 3 sessions; monkey 3: 7 sessions). Indeed, when reward is differently manipulated for go and stop trials, as for Go+ and Stop+ conditions in this task, the independence could be compromised as previously shown^17,38,39^.

We estimated the SSRT using the so-called integration method: for any given SSD (fixed-SSDs procedure), go RTs are rank ordered and the *n*^th^ go RT is selected, where *n* is the number of go RTs multiplied by the probability of responding at a given SSD. SSRT is obtained by subtracting the SSD from *n*^th^ RT. Since this method assumes the SSRT to be a constant value, averaging the SSRTs obtained at different SSDs provides the final SSRT estimation^38^. To compare SSRTs over the three cue conditions, we performed a Friedman test for repeated-measures (*p* < 0.05), testing together the SSRTs calculated from the different sessions of the three monkeys.

### Electrophysiological recordings and data analysis

Unfiltered raw activity was recorded from 96-channels Utah arrays (Blackrock Microsystems, USA) by using specific software (Tucker Davies Technologies, sampling rate 24.4 kHz). Array data in this paper come from one session for monkey 1, and two sessions for monkey 2, selected based on the best performance and adherence to the model. We had the possibility of using two sessions only in monkey 2 because they were sufficiently separated in time, to reduce oversampling from the same population of neurons (2.5 months interval between sessions).

We extracted the multiunit activity (MUA) by computing the time-varying power spectra P(ω,t) from the short-time Fourier transform of the recorded unfiltered electric field potential (UFPs) in ±40 ms sliding windows (20 ms steps). P(ω,t) was normalized by their average P_ref_(ω) across the first 1200 sec of the recording. Our spectral estimated MUAs were the average R(ω,t) across the 0.3-2.0 kHz band. The extracted MUA is considered a good approximation of the average firing rate, as described in detail in Mattia et al. ^40^.

All the analyses were conducted using Matlab software (The MathWorks Inc., Natic, MA, USA), and statistical comparisons were performed by means of Matlab and the software Statistica (StatSoft, Inc.).

### Task-related activity

We first selected array channels (recordings) based on their modulation during the task. We focused on three time periods: Cue-Early epoch (100-400 ms after the cue presentation), Cue-Late epoch (540-940 ms after the cue presentation) and Pre-MOV epoch (100-400 ms from Go signal). For each recording and each epoch, we compared the neuronal activity in correct go trials with the baseline period (from – 260 to −60 ms before the cue appearance) using a non-parametric Wilcoxon test (*p* < 0.001). We classified the neuronal activity as task-related whether activity during baseline was significantly different from activity in at least one of the three epochs. All the following analyses were accomplished on task-related recordings.

### Neuronal modulation related to the cue presentation on go trials

Task-related neuronal activities were examined to explore how they integrate information about the cue into motor preparation. To this end, we decided to concentrate on two time periods of the cue epoch (due to its duration = 1 sec): Cue-Early (+100 +400 ms from cue), and Cue-Late (+540 +940 ms from cue); and we compared them with the Pre-MOV epoch (+100 +400 ms from Go), i.e. the movement preparation epoch. We employed a one-way ANOVA to compare the neuronal activity over the different cue conditions, for each recording and each epoch. When *p* value was significant (*p* < 0.05), we used multiple comparison post-hoc tests to classify recordings, in each epoch of interest, based on the pattern of neuronal activity presented in the different cue conditions. We depicted three main categories that together described the neuronal modulation of most recordings: “Go+ & Stop+ > Neutral” included those recordings showing higher activity for Go+ and Stop+ compared to Neutral, and this difference was significant at least for one comparison (Go+ vs. Neutral or Stop+ vs. Neutral); “Go+ > Neutral & Stop+” classified recordings in which Go+ was higher than Neutral and Stop+, whereas these two were not different; lastly, recordings whose activity was higher in Go+ compared to Neutral and, in turn, Stop+ were included in the category “Go+ > Neutral > Stop+”. We then performed a chi square test (*p* < 0.05) on the frequencies of the three main categories across the three epochs of the task.

To assess how the neuronal activity changes across trial epochs based on the cue, we estimated two contrast indices (CI) between the two salient cue conditions versus the Neutral condition (i.e. *Go+ vs. Neutral*, and *Stop+ vs. Neutral*), both during Cue-Early and Pre-MOV epochs. CI were intended as:

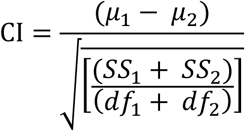

where μ_x_ is the mean neuronal activity, SS_x_ is the sum of squares, and df_x_ is the degree of freedom (number of trials minus 1) for each condition^41^. An index superior to zero indicated a stronger modulation in the salient cue conditions, while an index inferior to zero indicated a stronger modulation in the Neutral cue condition. We then performed a kruskal-wallis test to verify if contrast indexes significantly differed (*p* < 0.05) over time.

Finally, to test if the neuronal activity on different time epochs was explained by reward-related factors (REWARD), i.e., cues salience during Cue-Early epoch and motivation to move during Pre-MOV, when the effects of behavioural reaction times (RT) were factored out, we performed a multiple regression analysis, fitting three models:

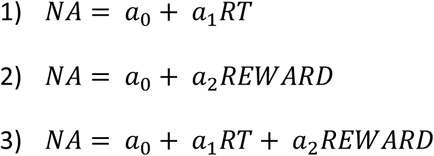

where NA is the mean neuronal activity measured during Cue-Early or during Pre-MOV epochs. The variable REWARD was set differently during the two epochs of the task: during Cue-Early we test the effect of cues salience, so attributing 1 to Go+ and Stop+ and 0 to Neutral trials; whereas, during Pre-MOV we test the influence of the motivation to move, assigning 2, 1 and 0 for Go+, Neutral and Stop+ trials, respectively. To determine whether adding the variable REWARD produced a significant improvement in performance, we compared *model 3* to *model 1*. To determine whether adding the variable RT produced a significant improvement, we compared *model 3* to *model 2*. Significance was assessed with an F-test using:

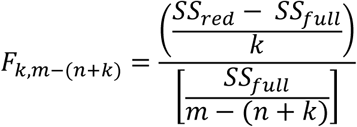

where k = 1 is the difference in degrees of freedom between the two models, *n* = 1 is the number of recordings, and m is the number of trials on which the analysis was based. SS_full_ and SS_red_ are the residual sums of squares resulting when the data were fitted with the full model (*model 3*) and the reduced model (*model 1* or *model 2*), respectively. The criterion for statistical significance was taken as *p* < 0.05 (for similar approach, see ^16^).

### Neuronal modulation related to the cue presentation on stop trials

To study the contribution of motivation to movement inhibition, we firstly selected recordings with neuronal activity involved in movement inhibition (countermanding or CMT recordings). In the framework of the stop-signal paradigm, we expect that CMT recordings to be differently modulated when the movement is inhibited respect to when the movement is made, and that the level of activity during correct stop trials changes early enough to be able to influence the cancelation of the movement (i.e. after the Stop signal presentation and before the end of the estimated SSRT). Against this background, we compared the neuronal activity on correct stop trials with that one on latency matched go trials. These are go trials in which RTs are longer than the specific SSD plus the corresponding SSRT, i.e. those trials that have a similar level of movement preparation of the correct stop trials. The analysis was accomplished for each SSD separately, putting together the different cue conditions and examining the time from the Go signal to 50 ms after the corresponding SSRT, in 20 ms time bins. We excluded the longest SSD (the fourth) from the analysis because of the few correct stop trials. A recording was classified as CMT if the ANOVA was significant (*p* < 0.01 for at least two consecutive bins after the Stop signal presentation) in at least two of three SSDs (for similar approach, see ^42^).

We then compared the neuronal activity on correct stop and latency matched go trials separately for each cue condition by means of receiver operating characteristic (ROC) analysis. For each CMT recording, we computed a measure of accuracy of the divergence of the neuronal activity (step 20 ms), starting 100 ms before the Stop signal to the end of SSRT. Some recordings (3 in monkey 1 and 2 in monkey 2) showed an opposite pattern of modulation compared to most of them: after the Stop signal their activity increased relative to the one observed in latency matched go trials (for a similar observation see ^12^). For these recordings we inverted ROC values (1 - ROC value) to compare them with the others. For recordings with significant activity difference (ROC value > 0.65 for at least 3 consecutive bins), we defined the onset of this difference as the first significant time bin. We refer to this time as the stop neuronal time (SNT).

We evaluated if the SNTs differed significantly based on the cue conditions performing a one-way ANOVA (*p* < 0.05) between the latency distributions. SNT estimations that occurred before or within 70 ms following the Stop signal presentation were excluded from this analysis. We also tested if the proportions of CMT recordings obtained by means of the ROC in the three cue conditions were different through z score tests (*p* < 0.05) estimated separately for each pair of conditions.

Lastly, we examined more in depth the neuronal dynamics of recordings modulated after the Stop signal presentation. To this aim we excluded those recordings (3 in monkey 1 and 2 in monkey 2) showing an increased activity in correct stop trials compared to latency matched go trials. We calculated the time from the Stop signal presentation at which the neuronal activity began to show a decreasing trend that went on for at least 100 consecutive ms. If this decreasing trend was not found, we excluded the corresponding recording from the following analysis. In total, 66, 75 and 41 recordings were analysed respectively for Go+, Neutral and Stop+ conditions. The time at which the neuronal activity began to show a decreasing trend was defined as the decrease onset time. We also examined the mean neuronal activity at the decrease onset time. Lastly, we considered 200 ms starting from the decrease onset time, and we ran a robust regression function to extract the slope of the neuronal suppression pattern. To test the influence of the motivational context over these inhibitory measures we performed three separate one-way ANOVAs (*p* < 0.05) - one for the decrease onset times, the other for the corresponding neuronal activity, and the last one for the slopes.

### Neural decoding analysis

We performed a neural population decoding analysis using the maximum correlation coefficient classifier previously described^43^. The classifier was trained to discriminate, among the different cue conditions organized on the basis either of the salience or of the motivation, and between the two possible correct responses of the stop-signal task - i.e. movement generation in correct go-trials and movement inhibition in correct-stop trials. We provided as input the multi-unit activity calculated in ±40 ms bins sampled at 20 ms intervals for each trial (data point), from each recording. For this population analysis, we combined the 80 task-related recordings from monkey 1, and the 31 task-related recordings from the session from monkey 2 with the highest number of recordings. Note that this analysis was performed on the entire population of task related recordings, also including non CMT recordings.

We defined the optimal split factor (k) as the highest number of trials available in each condition for each recording. Particularly, the decoding analyses of the “motivation” (Go+ vs. Neutral vs. Stop+ conditions, putting together correct go and correct stop trials) and “cue salience” (Go+ & Stop+ vs. Neutral conditions, putting together correct go and correct stop trials) factors were performed choosing a k of 164 (i.e., 164 trials x 3 conditions = 492 data points) and 219 (i.e., 219 trials x 2 conditions = 438 data points) respectively. Similarly, for the decoding analysis of the trial-type factor (correct go trials vs. correct stop trials), we used a k of 84 (i.e., 84 trials x 2 conditions = 168 data points), without having to remove any recording. The training of the classifier was then performed with pseudopopulations of recordings obtained by randomly arranging all the available data points into a number of split equal to the number of data points per condition, and normalized into z-scores to avoid that recordings with higher levels of activity dominated the decoding procedure. The classifier was trained using k – 1 number of splits and then tested on the remaining split: this procedure was repeated as many times as the number of splits. The overall decoding procedure was run 5 times with different selection of data in the training and test splits, and the decoding accuracy from these runs was then averaged.

To assess whether the classification accuracy was above chance, a permutation test was performed by randomly shuffling the attribution of the conditions to the different trials, before re-running the full clutter-decoding experiment. This procedure was repeated 5 times to obtain a null distribution of the decoding accuracies to be compared with the accuracy of the real decoding. The significance level was considered reached if the real decoding accuracies were greater than all the ones of the shuffle data in the null distribution for at least 5 consecutive significant bins.

Furthermore, we performed a cross-temporal decoding analysis by training the classifier at one point in time and testing its decoding performance at either the same or a different time point, we have been able to investigate the dynamic nature (i.e. no correlation between different time points, corresponding to a strong diagonal band in the plot) in time in an experiment) or static (i.e., strong correlation for different time points, corresponding to a strong square region in the plot) of the population code underlying the representation of “cue salience” and “motivation” in PMd.

## Availability of data and materials

The datasets used and/or data analysed during the current study are available from the corresponding authors on reasonable request.

## Authors’ contributions

PP and SF conceived the study; PP and FF designed the research; MG, FF, FG recorded the data; MG, SF, EB, PP, analysed and interpreted the data. MG, SF, PP wrote the paper. All authors critically commented and approved the final manuscript.

## Ethics approval and consent to participate

All experimental procedures conformed to European (Directive 2010/63/UE) and Italian (D.L. 26/2014) laws on the use of nonhuman primates in scientific research and were approved by the Italian Ministry of Health.

