## supplemental file for "Neuronal activity in the premotor cortex of monkeys reflects both cue salience and motivation for action generation and inhibition"

### SUPPLEMENTARY INFORMATION

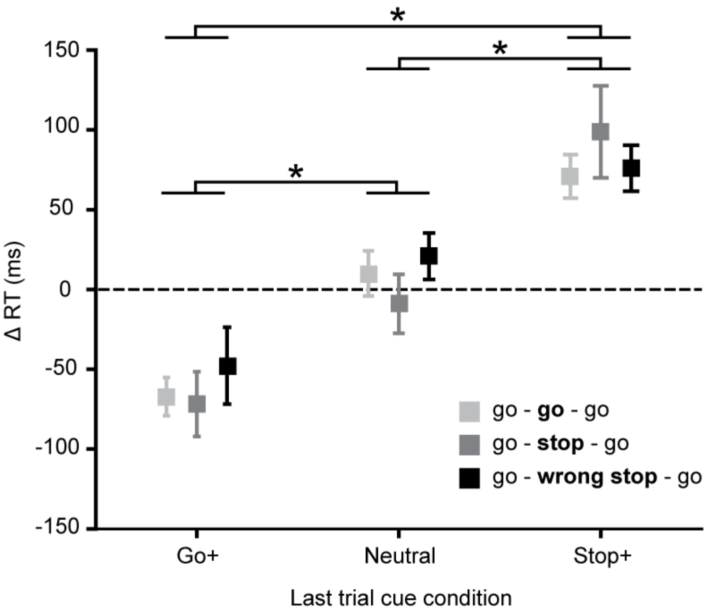


**Supplementary Figure 1.**

**Behavioural performance in the motivational stop-signal task in relation to trial history and cue condition.**

The differences between RTs (ΔRT) between correct go trials (last trial on each triplet; following correct go trials, correct stop trials and wrong stop trials) and the close in time go trial (first trial on each triplet) are shown (mean ΔRT ± SEM, n = 14 sessions) sorted by cue condition of the same last trials. A two-way ANOVA with as factors cue condition and trial sequence show that only the Factor cue condition was significantly different ($p < 0.05$; trial history $p = 0.701$) suggesting that animals were strongly influenced more by the context indicated by the cue than by the recent experienced trial type. The asterisk (*; $p < 0.05$) shows the results of post-docs (multiple comparison tests) between conditions.

**Methods related to Supplementary Figure 1.** We investigated whether the behavioural performance in our version of stop-signal task can be better explained by trial history (i.e., effect of the recent type of trial) rather than cue condition. Indeed, while performing the stop-signal task, subjects typically modulate their RTs based on the recent trial history (Emeric et al., 2007; Marcos et al., 2013; Montanari et al., 2017; Pouget et al., 2011). With this purpose, for each session (n = 14 sessions coming from all monkeys together), we sorted trials looking for triplets of successive trials: (go - go - go triplet) formed by the sequence *correct go trial, correct go trial, correct go trial*; (go - stop - go triplet) formed by the sequence *correct go trial, correct stop trial, correct go trial*; (go - wrong stop - go triplet) formed by the sequence *correct go trial, wrong stop trial, correct go trial*. For each of these sequences, we subtracted the RT of the first go trial from the RT of the last go trial, obtaining post-trial ΔRT. This procedure should help in providing effects related to the close critical trials (i.e., the central trial), avoiding global effects of RT variations (Dutilh et al., 2012; Nelson et al., 2010). The last component of each sequence was distinguished based on the cue condition to observe variations in this trial based on the cue compared to the preceding critical trial.

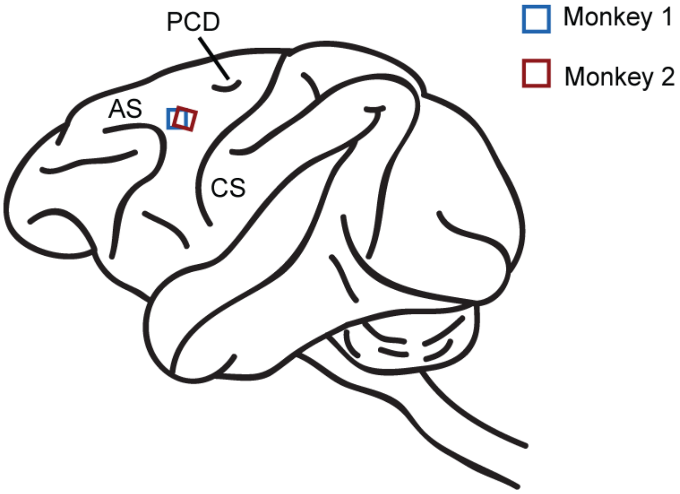


**Supplementary Figure 2.**

**Array locations on the dorsal premotor cortex (PMd) of monkey 1 and monkey 2.** Recordings were performed in the hemisphere contralateral to the arm used during the experiment. This illustrative brain figure represents the locations of arrays based on reconstruction from photos taken during surgery after dura opening and array insertion. AS: arcuate sulcus; PCD: pre-central dimple; CS: central sulcus.

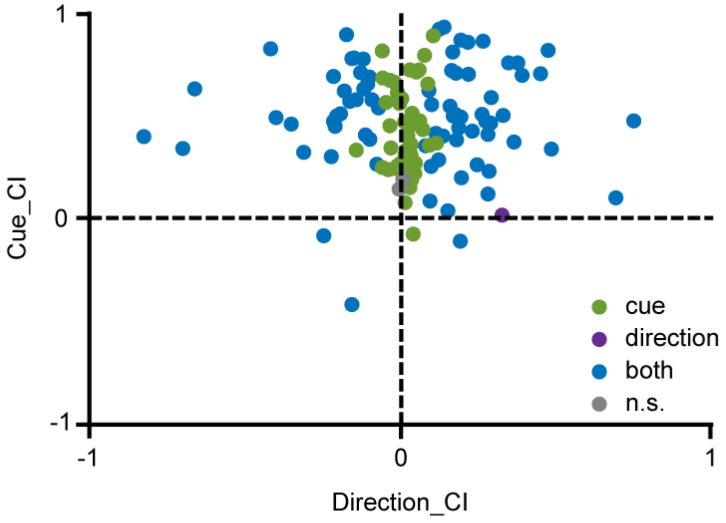


**Supplementary Figure 3.**

**Effect of cue condition and movement direction on the neuronal activity during go trials.** Scatter plot of cue-versus direction-contrast indices for each recording (n = 124 recordings) calculated during Pre-MOV epoch (100-400 ms after Go signal). Cue-contrast indices (Cue_CI) >0 indicate higher activity for the Go+ condition, and indices <0 indicate higher activity for the Stop+ condition. Direction-contrast indices (Direction_CI) >0 indicate higher activity when the movement is performed on the right; values <0 indicate higher activity when the movement is performed on the left. Symbol colour indicates those recordings with a significant main effect (two-way ANOVA, $p < 0.05$) of cue condition (green), movement direction (purple), or both (blue). Grey symbols represent those recordings with a not significant main effect (n.s.).

**Methods related to Supplementary Figure 3.** To quantify the influence of cue conditions and movement directions on movement-related activity, we estimated for each recording (124 recordings) a cue-contrast index (Go+ vs. Stop+) and a direction-contrast index (right vs. left) during the Pre-MOV epoch (100-400 ms after Go signal). Contrast indices (CI) were intended as in the main text (see Methods).

We additionally performed a two-way ANOVA (with cue condition and movement direction as factors; $p < 0.05$) to classify recordings whose neuronal activity was better explained by cue condition, movement direction or both.

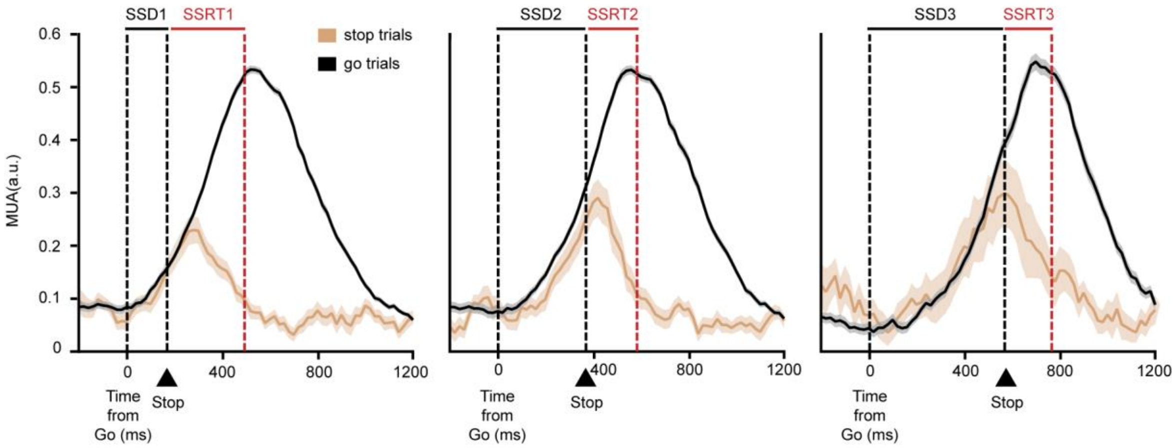


**Supplementary Figure 4.**

**Example of countermanding (CMT) recordings.** Example from monkey 1. Left, central and right panels represent the neuronal activity modulation for trials with different stop signal delays (SSDs) - SSD1, SSD2 and SSD3, respectively. Each panel shows the average MUA (± SE) of correct stop trials (stop trials; brown lines) and latency matched go trials (go trials; black lines). The three dashed lines represent the Go signal, the Stop signal and the end of the SSRT, respectively. SSDs and corresponding SSRTs durations are depicted at the top of each panel. For all, the difference in the neuronal activity starts after the Stop signal presentation and before the end of the estimated stop signal reaction time (SSRT).

**Supplementary Figure 5.**

**Neural decoding applied to trial-type.**

Classification accuracy over time and cross-temporal decoding plot for the trial-type (correct go trials vs. correct stop trials). The superimposed white line represents the classification accuracy of the population (n = 111 recordings). The red line in the lower part of the plot indicates the period of time during which the classification accuracy is significantly above chance level (chance level is 50%). White dashed lines indicate the time when the cue and the go signal were presented. Cross-temporal decoding plot has been obtained following the same procedure as in main Fig.5.

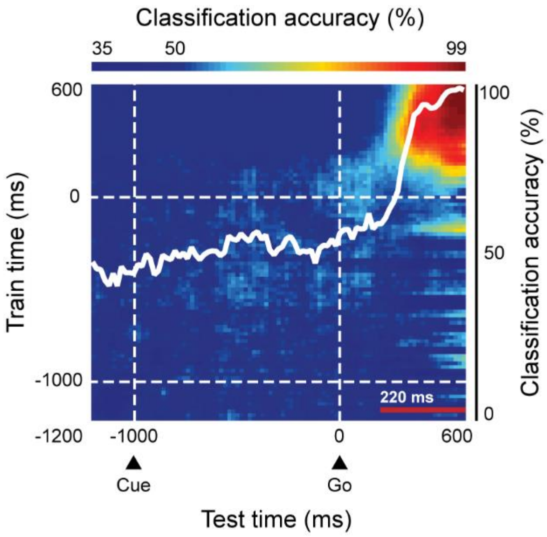

| ***Cue condition*** | | ***Go+*** | ***Neutral*** | ***Stop+*** |
|---|---|---|---|---|
| ***Go trials*** | | | | |
| Error rate ± SE (%) | Monkey 1 | 7 ± 0 | 6 ± 3 | 59 ± 17 |
| | Monkey 2 | 11 ± 0 | 24 ± 2 | 41 ± 10 |
| | Monkey 3 | 3 ± 0 | 6 ± 1 | 10 ± 3 |
| Mean RT ± SE (ms) | Monkey 1 | 593 ± 4 | 675 ± 5 | 786 ± 11 |
| | Monkey 2 | 480 ± 6 | 688 ± 7 | 731 ± 9 |
| | Monkey 3 | 599 ± 2 | 619 ± 3 | 692 ± 4 |
| ***Stop trials*** | | | | |
| Error rate ± SE (%) | Monkey 1 | 69 ± 3 | 50 ± 8 | 23 ± 10 |
| | Monkey 2 | 84 ± 2 | 46 ± 5 | 22 ± 6 |
| | Monkey 3 | 62 ± 1 | 47 ± 2 | 28 ± 2 |
| SSRT ± SE (ms) | Monkey 1 | 269 ± 8 | 250 ± 30 | 128 ± 110 |
| | Monkey 2 | 157 ± 17 | 176 ± 40 | 69 ± 37 |
| | Monkey 3 | 196 ± 30 | 159 ± 19 | 146 ± 13 |

**Supplementary Table 1.** Behavioural measures calculated separately from each monkey.
